# Disentangling spatial and environmental effects: flexible methods for community ecology and macroecology

**DOI:** 10.1101/871251

**Authors:** Duarte S. Viana, Petr Keil, Alienor Jeliazkov

## Abstract

Community ecologists and macroecologists have long sought to evaluate the importance of environmental conditions in determining species distributions, community composition, and diversity across sites. Different methods have been used to estimate species-environment relationships, but their differences to jointly fit and disentangle spatial autocorrelation and structure remain poorly studied. We compared how methods in four broad families of statistical models estimated the contribution of the environment and space to variation in species binary occurrence and abundance. These methods included distance-based regression, generalized linear models (GLM, and the special case of RDA), generalized additive models (GAM), and tree-based machine learning (ML): regression trees, boosted regression trees (BRT), and random forests. The spatial component of the model consisted of spatial distance (in distance-based regression), Moran’s Eigenvector Maps (MEM; in GLM and ML), smooth spatial splines (in GAM), or tree-based non-linear modelling of spatial coordinates (in ML). We simulated typical site-by-species data to assess the methods’ performance in (1) fitting environmental and spatial models, and (2) partitioning the variation explained by environmental and spatial predictors. We observed marked differences in performance mostly caused by imbalanced performance in estimating environmental and spatial effects. Such differences also manifested when analyzing eight different empirical datasets. GLM and BRT with MEMs were generally the most reliable methods for partitioning the variation explained by environmental and spatial effects across a wide range of simulated scenarios. The remaining methods tended to underfit simulated spatial structures, causing underestimation of spatial fractions of variation. Our results suggest that previously overlooked methods for performing variation partitioning, especially tree-based ML, offer flexible approaches to analyze site-by-species matrices. We provide general guidelines on the usefulness of different models under different ecological and sampling scenarios, for species distribution modelling, community ecology, and macroecology.

## Introduction

The environment is a major driver of species occurrence and abundance, shaping many facets of biodiversity, from fine-scale community composition to large-scale species distributions and co-occurrence (Chase and Leibold 2003, Townsend Peterson et al. 2011). Consequently, the environment is central in ecological theory, including coexistence theory (Chesson and Warner 1981, Chesson 2000), modern niche theory (Chase and Leibold 2003), metacommunity theory (Leibold et al. 2004, Thompson et al. 2020), as well as biogeographical and macroecological theory (Ricklefs and Jenkins 2011, Townsend Peterson et al. 2011). Species-environment relationships (SERs) have been widely estimated to (i) characterize the species’ niches and model species distributions and community composition (Townsend Peterson et al. 2011, Bar-Massada 2015, Norberg et al. 2019), (ii) explore the importance of environmental filtering as a biodiversity process (Cottenie 2005, Soininen 2014), and (iii) correct for environmental effects when studying biotic interactions and other community assembly processes (e.g. Ovaskainen et al. 2017, D’Amen et al. 2018).

One of the major challenges when studying SERs is the correct estimation of environmental effects while accounting for spatial autocorrelation of species distributions at different spatial scales, caused by both environmental autocorrelation and spatial processes such as dispersal limitation. In particular, a popular aim in community ecology has been to disentangle, using variation partitioning, the relative importance of environmentally driven (niche) processes from spatial processes often associated with neutral theory (Cottenie 2005, Peres-Neto et al. 2006, Soininen 2014, Leibold and Chase 2017).

However, estimating SERs while accounting for spatial autocorrelation and/or performing variation partitioning has been done differently in different research fields. In biogeography and macroecology, generalized linear and additive models (GLM and GAM) as well as tree-based machine learning (ML) methods are widely used to estimate SERs (e.g. Elith and Graham 2009, Norberg et al. 2019). In community ecology multivariate methods, in particular constrained ordination (such as CCA and RDA) or distance-based methods, have been widely used to partition explained variation in community composition according to environmental and spatial effects and infer community assembly processes (Cottenie 2005, Peres-Neto et al. 2006, Tuomisto and Ruokolainen 2006, Soininen 2014). However, these methods have been criticized, since their performance depends on the strength of effects, the spatial structure of the environmental variables, and model specification (e.g. Gilbert and Bennett 2010, Smith and Lundholm 2010).

On the other hand, methods used in biogeography and macroecology such as GLM, GAM, and ML, have been widely overlooked for disentangling spatial dependence in the context of variation partitioning, particularly in a community context with multi-species responses. Machine learning methods such as Random Forest and Boosted Regression Trees (Elith and Graham 2009), and their multivariate versions (Nieto-Lugilde et al. 2018), have several advantages over classical regression methods, particularly because they are usually bound by fewer statistical assumptions and are inherently suited to model complex interactions and non-linear relationships. Although comparisons of different models to estimate species distributions exist (Norberg et al. 2019), the usefulness of methods such as GAM and machine learning to model different spatial structures while estimating environmental effects, and to partition variation, has been less explored. Here, we compare the virtues and drawbacks of methods based on generalized linear models, generalized additive models, and machine learning (Table 1) to model and disentangle environmental and spatial effects. We do this comparison by using both simulated and empirical data. We specifically focus on site-by-species matrices of occurrence or abundance as response variables, but this study also applies to any other kind of response variable affected by spatial autocorrelation (e.g., species richness and beta-diversity).

**Table 1.**
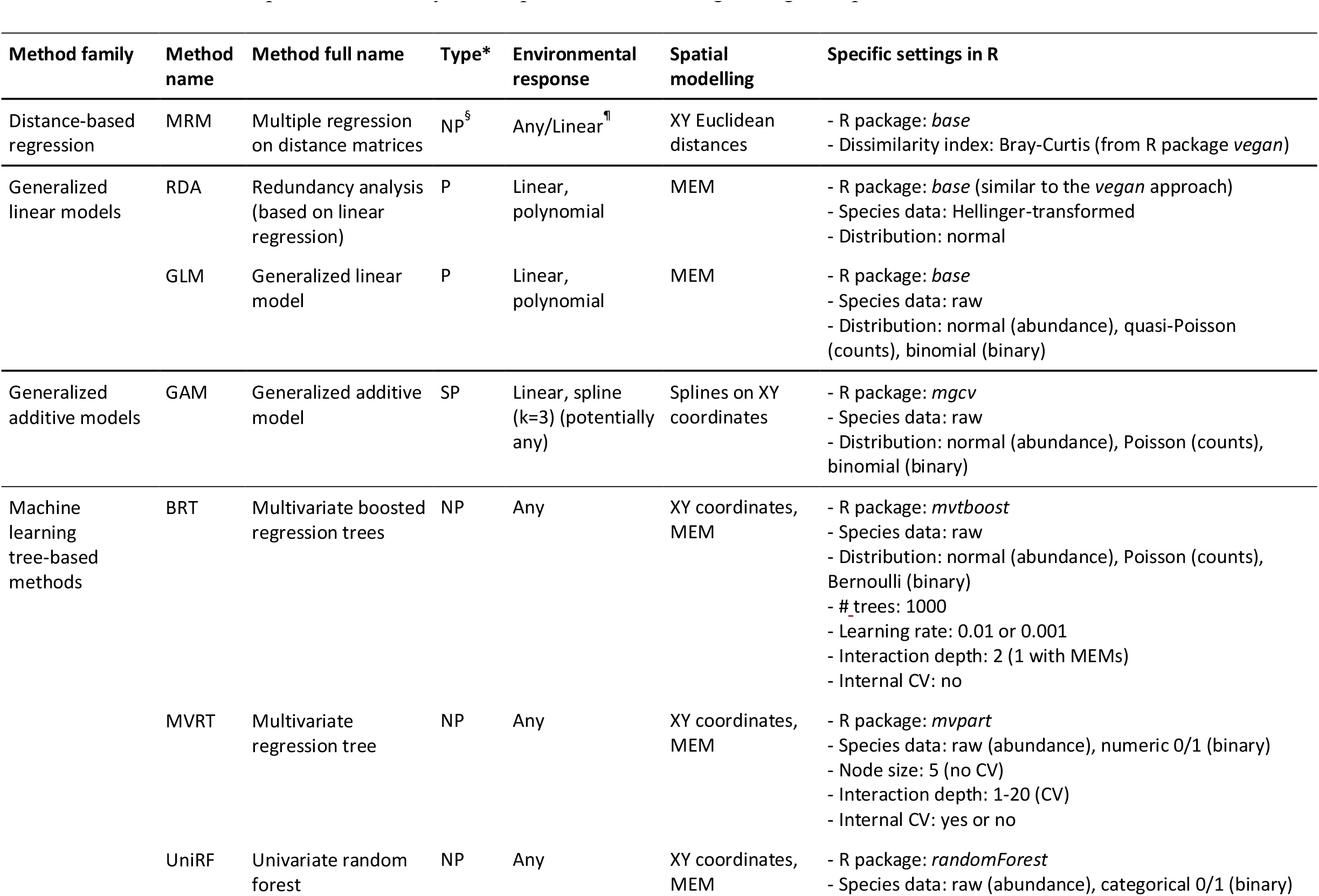
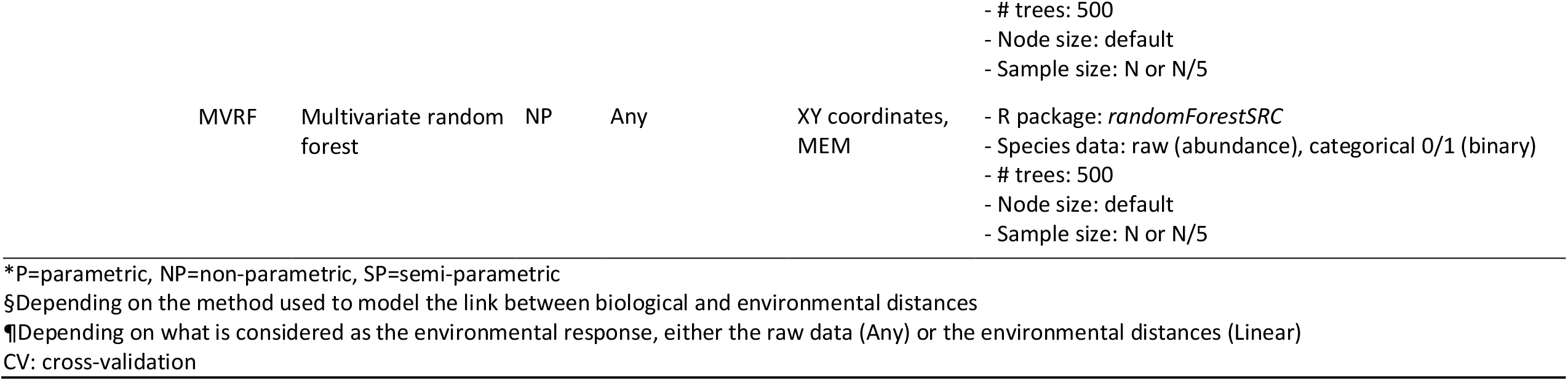
The methods compared in this study and respective model fitting settings and parameters.

## Methods

All the analyses were carried out in R (R Development Core Team 2020), and all the computer code is available at https://github.com/duarte-viana/iVarPart.

### Data simulation

We simulated site-by-species community data where variation in abundance or binary occurrence corresponded to pre-defined species’ responses to environmental conditions, spatial gradients, or both. We used a simple simulation consisting of a grid of *N* cells (hereafter sites), where each site was occupied by a different community and was environmentally homogeneous. The occurrence or abundance 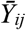 of each species *i* at a given site *j* depended on an environmental response (*X*_*ij*_), a spatial response (*W*_*ij*_), or their linear combination.

#### Environmental response (X)

We simulated one spatially autocorrelated environmental variable *E* (0 *≤ E ≤* 1) on the grid by simulating a random Gaussian field in which the autocorrelation level was set by the range parameter (*A*) of an exponential variogram model (here fixed at *A*=*N*/2), using the R package *gstat* (Pebesma 2004). Species responded to *E* either linearly or according to a Gaussian curve (i.e. a unimodal, bell-shaped response). As such, the response to the environment (*X*_*ij*_) of species *i* in cell *j* was

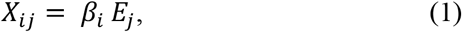

where *E*_*j*_ is the environmental value in cell *j* and *β*_*i*_ is the slope of the linear response (*β*_*i*_ was taken randomly from a normal distribution of *β* values; mean=10, s.d.=2). Alternatively,

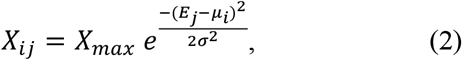

where *μ*_*i*_ is the mean of the Gaussian response (i.e. the optimal environmental condition for species *i, μ* being equally spaced along the environmental variable from 0.05 to 0.95), *σ* is the standard deviation (i.e. the fundamental niche breadth), and *X*_*max*_ (=20) is a constant defining the maximum abundance. For examples of the linear and Gaussian-shaped responses, see Appendix S1: Figure S1.1.

#### Spatial response (W)

The spatial response *W* was defined by simulating another random Gaussian field *S* on the same grid, independently from *E*, and setting the abundance (or occurrence) *W*_*ij*_ to be directly proportional to *S*_*ij*_ (*W*_*ij*_ *= X*_*max*_ *S*_*ij*_). *X*_*max*_ (=20) sets *W*_*ij*_ to the same scale as *X*_*ij*_. We simulated different types of spatial structures *S* by defining two different spatial models (an exponential or Gaussian variogram model) with varying ranges (*A*). For examples of spatial patterns of *S*, and thus *W*, see Appendix S1: Figure S1.2.

#### Multi-species responses 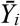

In order to vary the relative contributions of the environment (*X*) and space (*W*) to variation in species’ abundance or occurrence, *X*_*i*_ and *W*_*i*_ were given different weights (*β*_*X*_ and *β*_*W*_, respectively; *β*_*X*_ + *β*_*W*_ = 1), so that

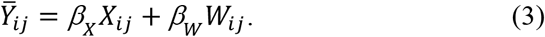

The simulated site-by-species matrix was obtained as follows. Occurrence (i.e. presence-absence) data *Y*_*ij*_ were obtained by first transforming 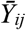(centered around 0) into probabilities *P*_*ij*_ using an inverse logit function, and then randomly sampling from a Bernoulli distribution with parameter *P*_*i*_. Two types of abundance data *Y*_*ij*_ were generated: data with normally distributed errors (*μ* = 0 and *σ* = 2) and data with Poisson distributed errors using the predicted abundances 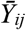as the mean abundances of the Poisson distribution (see an example of the spatial pattern of the resulting *Y* in Appendix S1: Figure S1.3). The relative contribution of *X* and *W* to variation in *Y* was measured by partitioning the squared correlation coefficient *r*^*2*^ between *Y*_*i*_ and 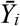(equivalent to *R*^*2*^ in a linear regression) into the total fraction [*X*] of variation attributable to *X* and the pure fraction [*W*] of variation attributable to *W*, according to:

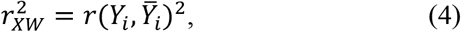

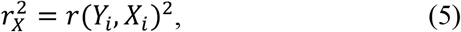

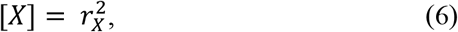

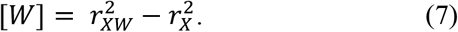

Although *W* is independent from and thus uncorrelated with *X*, some spurious collinearity between them can arise and jointly explain variation in *Y*. Because *W* is a pure spatial variable, the collinear effect is the effect of spatially correlated *X*. Note that fraction [X] contains this shared fraction of variation, unlike the pure fraction of variation [W]. As expected, *β* was directly related to the partial correlation *r* between *X*_*i*_ or *W*_*i*_ and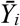, i.e., 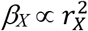and 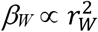 (Appendix S1: Figure S1.4). The *r*^*2*^ values were Pearson correlation for normal data, Spearman rank correlation for counts data, and point-biserial correlation (as defined in the R package *ltm*; Rizopoulos 2006) for binary data.

We considered different scenarios in the type of data (normal, counts, or binary), the size of the grid (*N* = 25, 100, or 400 sites), the type of spatial variogram model (exponential or Gaussian), the autocorrelation range of *W* (*A* = 0.01*N*, 0.5*N*, or *N*), and the type of response to the environment, either linear or bell-shaped with varying niche breadth (*2σ*^*2*^ = 0.002, 0.02, or 0.2) (see also Appendix S1: Table S1). The reference scenario while one target parameter varied was *N* = 100, *A* = 0.5*N* and 2*σ*^2^ = 0.02. In addition, we also considered a scenario where a random subsample (50 out of 400 sites) was taken to simulate a sampling effect, and another scenario where either 3 or 6 random environmental variables orthogonal to the response (i.e. noise) were added to inspect overfitting propensity. For each combination of simulation parameters, we simulated a range of predictive weights *β*_*X*_ from 0 to 1 increasing by 0.1, determining the relative importance of environment and space, since *β*_*W*_ =1-*β*_*X*._ Each combination of parameters and *β*_*X*_ was replicated 5 times, resulting in a total of 3465 simulation runs.

### Statistical methods

We considered methods in four broad families of statistical models (Table 1): distance-based regression, generalized linear models (GLM), generalized additive models (GAM), and tree-based machine learning (ML). These methods can fit species-environment and species-space relationships in a variety of ways as different environmental and spatial predictors can be used to estimate the same environmental and spatial effects. Because considering all the possible approaches with each method would result in an intractable list, we opted for the most widely used ones, as described below. Our goal was to assess the performance of these different approaches for fitting environmental and spatial effects (Exercise 1), and partitioning explained variation (Exercise 2).

#### 1. Distance-based regression

Applying regression on compositional distances between sites derived from site-by-species abundance or occurrence (i.e. pairwise beta-diversity) (Tuomisto and Ruokolainen 2006) has often been used to investigate the relative importance of environment and space in determining community assembly patterns (see Tuomisto and Ruokolainen, 2008, and the discussion initiated with Legendre et al. 2005; Weinstein et al., 2014; Robroek et al., 2017). The most common distance-based regression method is the Multiple Regression on distance Matrices (MRM), which is the adaptation of the Mantel regression to multiple regression analysis (Lichstein 2007). It consists of regressing ecological distances (i.e. compositional dissimilarities between sites; the response matrix) against the matrices of environmental and geographical distances (the predictor matrices). We used Bray-Curtis dissimilarities for the species data (R package *vegan*; Oksanen et al. 2017), and Euclidean distances for both the environment and the spatial coordinates. A simple linear regression was performed on the pairwise distances using function ‘lm’ of the R *stats* package (R Development Core Team 2020).

#### 2. Generalized linear models

We fitted GLMs using the *stats* R package (R Development Core Team 2020). We fitted either a linear effect or a second-degree polynomial to model the environmental effect (depending on the simulated environmental component *X*). To model the spatial effects, we used distance-based Moran’s Eigenvector Maps (MEM), calculated using the function “dbmem” (with default settings) of the R package *adespatial* (Dray et al. 2016). MEM variables represent the spatial autocorrelation across sampled sites at different spatial scales (i.e., different grains of autocorrelation). Also more broadly known as Principal Coordinates of Neighbourhood Matrices (PCNM), MEMs are widely used to model spatial structure and autocorrelation. All GLMs were fitted as single models to each species and the predictions were then stacked into a matrix of predicted species abundances or occurrences. The error distribution of the model was defined according to the type of simulated data: a normal distribution for normal errors, quasi-Poisson for counts data (due to best estimation convergence compared to Poisson and negative binomial) and binomial/Bernoulli distribution for binary data. We also included the linear models underlying RDA (van den Wollenberg 1977) in our comparison. The difference to GLM models is that the species data are usually first transformed and subsequently used to estimate a linear model for each species – note, therefore, that the error distribution is always assumed to be normal, independently of the type of data. Accordingly, prior to GLM, the response data were Hellinger-transformed (as recommended in Legendre and Gallagher 2001, Legendre and Legendre 2012) and, when the response to the environment was bell-shaped, log-transformed to improve model fitting.

#### 3. Generalized additive models

GAMs were fitted using the R package *mgcv* (Wood 2017). We fitted either a linear effect or a thin-plate spline with *k*=3 dimensions (for the case of unimodal responses) to model the environmental effect, and 2D thin-plate splines with either a default *k* (that depends on the number of predictors) or fixed *k*=10 on the spatial coordinates of the sites to model the spatial effect. The different *k* allowed us to compare different initial degrees of “wiggliness” of the basis functions. The smoothing parameters were estimated by restricted maximum likelihood (REML). Similarly to GLM, the error distributions were tailored to the type of data, but unlike in GLM, the Poisson distribution was preferred for counts data due to good convergence.

#### 4. Machine learning: tree-based methods

Tree-based methods (Hastie et al. 2009) recursively split the response along a set of predictor variables, resulting in one tree or multiple trees (i.e. a “forest”). In contrast to the previous methods, tree-based machine learning makes no assumption regarding the functional form of the SERs; instead, the relationship is learned from the data. As such, the environmental predictor was *E* and the spatial predictors were either the spatial coordinates (XY) or MEMs (see model specifications and settings in Table 1). When spatial coordinates were used, we tried two variants of the method by defining a different value for the algorithm parameter that mostly affected the smoothness of the fitted model, according to *a priori* trial-error explorations.

##### Multivariate Regression Trees (MVRT)

This method fits a single multivariate regression tree (De’ath 2002) to explain the abundance or occurrence data. We used its implementation in R function ‘mvpart’ (package *mvpart*). We fitted a tree with the minimum number of observations in any terminal (“leaf”) node fixed to 5. We fitted the models with or without internal cross-validation. The model with MEMs was estimated without cross-validation.

##### Univariate Random Forest (UniRF)

This fits a univariate random forest (Breiman 2001, Hastie et al. 2009) to each species individually, using ‘randomForest’ R function (package *randomForest*). A random forest consists of a set of regression trees fitted to bootstrapped (i.e. sampled with replacement) data, each tree fitted to a random fraction of predictors. Predictions of the individual trees are then averaged to get the overall prediction. The UniRF method is identical to the method called Gradient Forest implemented in package *gradientForest* (Ellis et al. 2012). We used the default settings of the ‘randomForest’ function, as we expect the defaults to be most often adopted by users – specifically, the random forest consists of 500 regression trees, each with a sample size of either *N* (the sample size of the data) or *N*/5 (i.e. 20% of the data), each fitted to data resampled randomly with replacement, using a random subset (1/3) of the predictors. The sample size was observed to influence the smoothness of the fitted model, thus being tightly related to the control of overfitting. The model with MEMs used a sample size of *N*.

##### Multivariate Random Forest (MVRF)

Similarly to the univariate version, this “multivariate” version fits a regression tree to each species, but now all happens within a single function call, and the split rule is the composite normalized mean-squared error (CNMSE), where each component (species) of the composite is normalized so that the mean abundance of the species does not influence the split rule. We used the R function ‘rfsrc’ (package *randomForestSRC*; Ishwaran et al. 2019). We used the minimum number of observations in any terminal (“leaf”) node fixed to 5, and followed the default settings of the R function so that each tree in the forest was fitted to data resampled randomly with replacement, and the number of randomly chosen predictors in each tree was sqrt(*p*), rounded up, where *p* is the total number of predictors. Such as for UniRF, we varied the sample size to be either *N* (the sample size of the data) or *N*/5 (i.e. 20% of the data) in order to assess different “smoothness” settings. Here again, the model with MEMs used a sample size of *N*.

##### Boosted Regression Trees (BRT)

This method uses a gradient boosting algorithm of Friedman (2001) which fits, to each species separately, a sequence of regression trees, where each new tree is applied to the residuals from the previous tree (Miller et al. 2016). We used the R function ‘mvtb’ (package *mvtboost*) and fitted two variants of the algorithm – shrinkage (or “learning rate”) set to either 0.01 or 0.1. We set the tree depth to two to allow the model to fit complex spatial structures resulting from the interaction between the spatial coordinates (x, y), except for the model with MEMs, where we used a tree depth of 1 to avoid modelling unnecessary interactions between MEMs. The model with MEMs used a sample learning rate of 0.01.

### Assessment of method performance on simulated data

Our study comprised two exercises. In Exercise 1 we assessed the flexibility of the methods in Table 1 to approximate the simulated linear or bell-shaped responses of species to the environment, as well as their responses to spatial structures which are typically more complex and non-linear. In Exercise 2 we partitioned the variation in multi-species abundance or occurrence caused by the spatial and environmental effects, and compared these to the simulated spatial and environmental fractions of explained variation.

#### Exercise 1. Model fitting performance

We applied each of the methods (Table 1) on simulated data and obtained the model predictions (*Ŷ*). The fitting performance was calculated by the correlation between the simulated abundance or occurrence values before adding the error structure (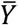; i.e. the true model) with the values predicted by the method *Ŷ*. However, deviations from the true model might be due to under- or overfitting: if data are overfitted, the predictions will be closer to the observed data *Y* (as the added error will be also fitted to some extent) and the correlation between *Y* and *Ŷ* will be higher than the correlation between 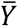 and *Ŷ*; if data are underfitted, the correlation between *Y* and *Ŷ*will be lower than that between 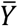 and *Ŷ*. We explicitly assessed these causes of underperformance. The performance was assessed for models estimating environmental and spatial effects jointly, environmental effects alone (i.e., when *β*_*X*_ = 1 and *β*_*W*_ = 0), and spatial effects alone (i.e., when *β*_*X*_= 0 and *β*_*W*_ = 1). RDA and MRM were excluded from this exercise, because the transformations of the response data make these models incomparable with the others.

#### Exercise 2. Variation partitioning performance

We performed variation partitioning with each of the methods according to the same procedure as for the simulated data (equations 4-7), but now we used the model predictions to calculate the *r*^*2*^ and obtain the fractions of explained variation 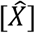and [*Ŵ*] Note that the spatial predictors in the statistical models also estimate the effect of the environment that is spatially autocorrelated, thus 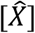also contains the variation attributed to the effect of spatially correlated environment. As such, the shared fraction of variation estimated from the statistical models was systematically higher than the shared fraction of the simulated data. This is why we compare the total (pure + shared) environmental fraction of variation. The performance of each method under the different simulated scenarios was assessed by comparing the estimated vs. simulated fractions of variation. To keep the comparison fair, we also used squared correlation coefficients (*r*^*2*^) for the estimated variation fractions. To check possible effects of the choice of goodness-of-fit metric, we also performed variation partitioning using *R*^*2*^ or deviance-based pseudo-*R*^*2*^ metrics depending on the type of data and method. For GLM methods (including RDA) we adjusted the *r*^*2*^ to account for the number of predictors (Peres-Neto et al. 2006). Negative fractions of variation were set to 0.

### Comparison of methods using empirical data

To explore the impacts of method choice on empirical results, we compared the different methods (Table 1) by performing variation partitioning on nine empirical datasets (see Appendix S2). Each empirical dataset consisted of a site-by-species abundance matrix, some environmental variables and geographical coordinates of the sites. For GLMs we included both linear and quadratic effects for the environmental predictors. Because the sample size of these data tended to be small (29<N<138), we limited the number of environmental predictors by randomly choosing 3 continuous predictors that were the same across the different models.

## Results

### Exercise 1. Model fitting performance

We observed clear differences in fitting performance among the different methods (Fig. 1). Most methods tended to underfit the data when both environmental and spatial effects were included, except GLM, BRT, and the MVRF variant with lower smoothness (MVRF-SS0 and MVRF-MEM) (Fig. 1a-c). The environmental effects were generally better fitted than the spatial effects, as the predictions match the true (simulated) response more closely (Fig. 1d-i). In particular, parametric or semi-parametric methods (GLM, GAM) showed the best fitting performance for the environmental effects (Fig.1d-f), except for data with normal error distribution because these models failed to accurately fit the bell-shaped response, even if the response variable was log-transformed (Appendix S1, Fig. S1.5b). Methods using MEMs tended to misfit the spatial component by either overfitting (GLM, BRT, MVRF) or underfitting it (UniRF, MVRT), but in general to a lesser extent than the other methods. The spatial splines in GAM and non-linear modelling of spatial coordinates in ML tended to underfit the spatial component, except BRT with the higher learning rate (Fig.1g-i).

**Figure 1.**
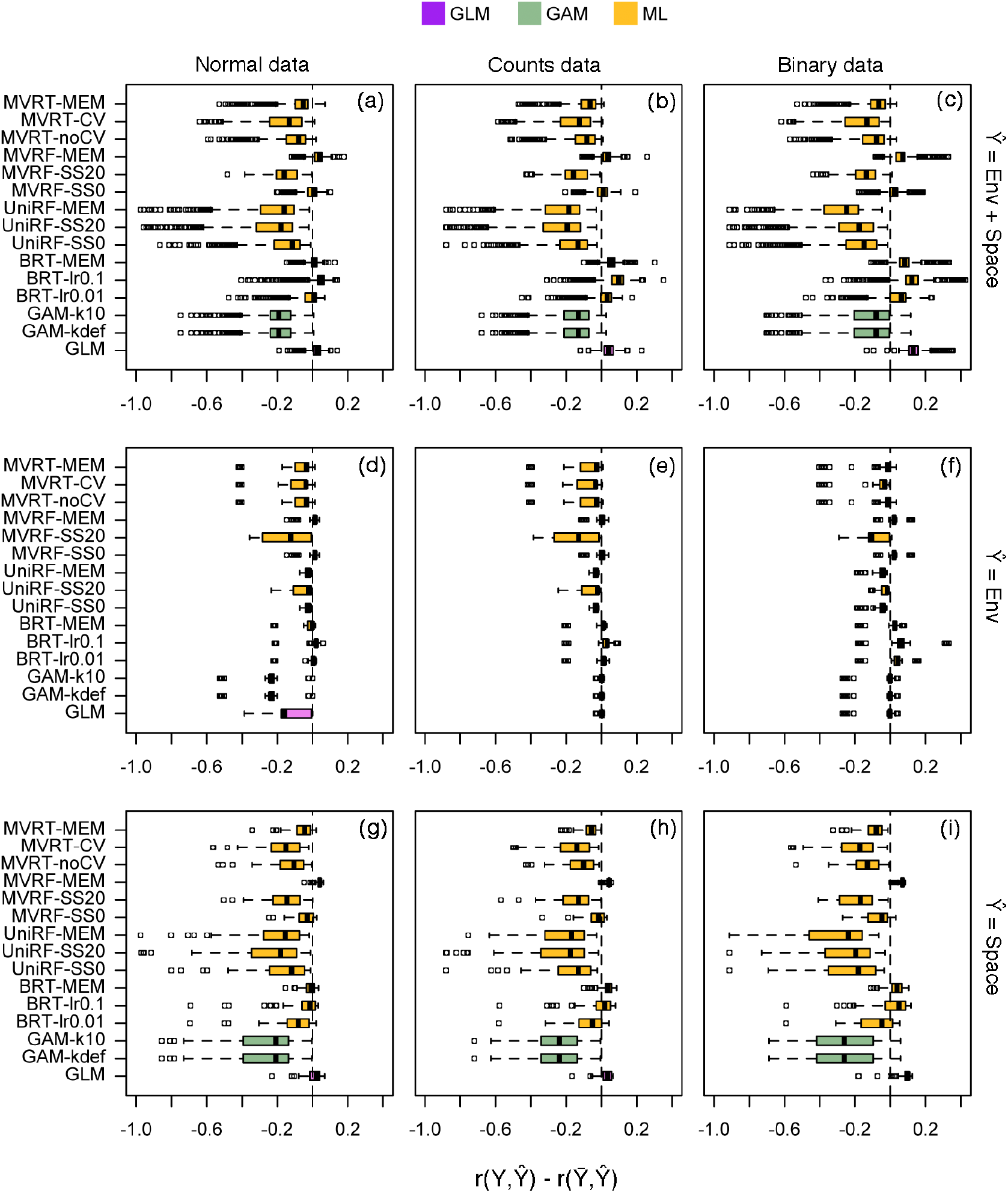
Fitting performance of the different statistical models for the additive effect of environment and space (a-c), the environmental effect alone (d-f), and the spatial effect alone (g-i), fitted to normal, counts, and binary data, across the different generated data (see Appendix S1: Table S1). The performance is given by the correlation between the observed response values (*Y*) and the model predictions (*Ŷ*) minus the correlation between the simulated response values 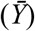 and the model predictions (*Ŷ*). The dashed line represents a perfect fit 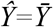, positive values indicate overfitting, and negative values indicate underfitting. MEM: Moran’s Eigenvector Maps, k: dimension of the spline basis function (default, “def”, or with fixed k=10, “k10”), lr: learning rate (0.01 or 0.1), SS: resample size (0 for SS=N and 20 for SS=N/5), CV: internal cross-validation (with, “CV”, or without, “noCV”).

The type of environmental response and spatial structure also caused fitting differences (Fig. 2). The parametric and semi-parametric methods (GLM and GAM) showed decreasing fitting performance as the shape of the bell-shaped environmental response narrowed, whereas BRT (and the other ML methods) could better fit these types of response, but could also easily overfit if not sufficiently smoothened. For the spatial effects, the fitting performance increases with the smoothness of the spatial pattern (Fig. 2). For a nearly random spatial structure (*A*=0.01*N*), for which we expected that the models did not fit the data (i.e. we expected simulated-predicted correlations to be close to 0, and thus a score closer to −1 in Fig. 2d, e, f), we observed that MEMs overfitted the data, BRT overfitted the data to some extent (unless learning rate was reduced), whereas GAM correctly did not overfit into this random variation (Fig. 2, lower panels).

**Figure 2.**
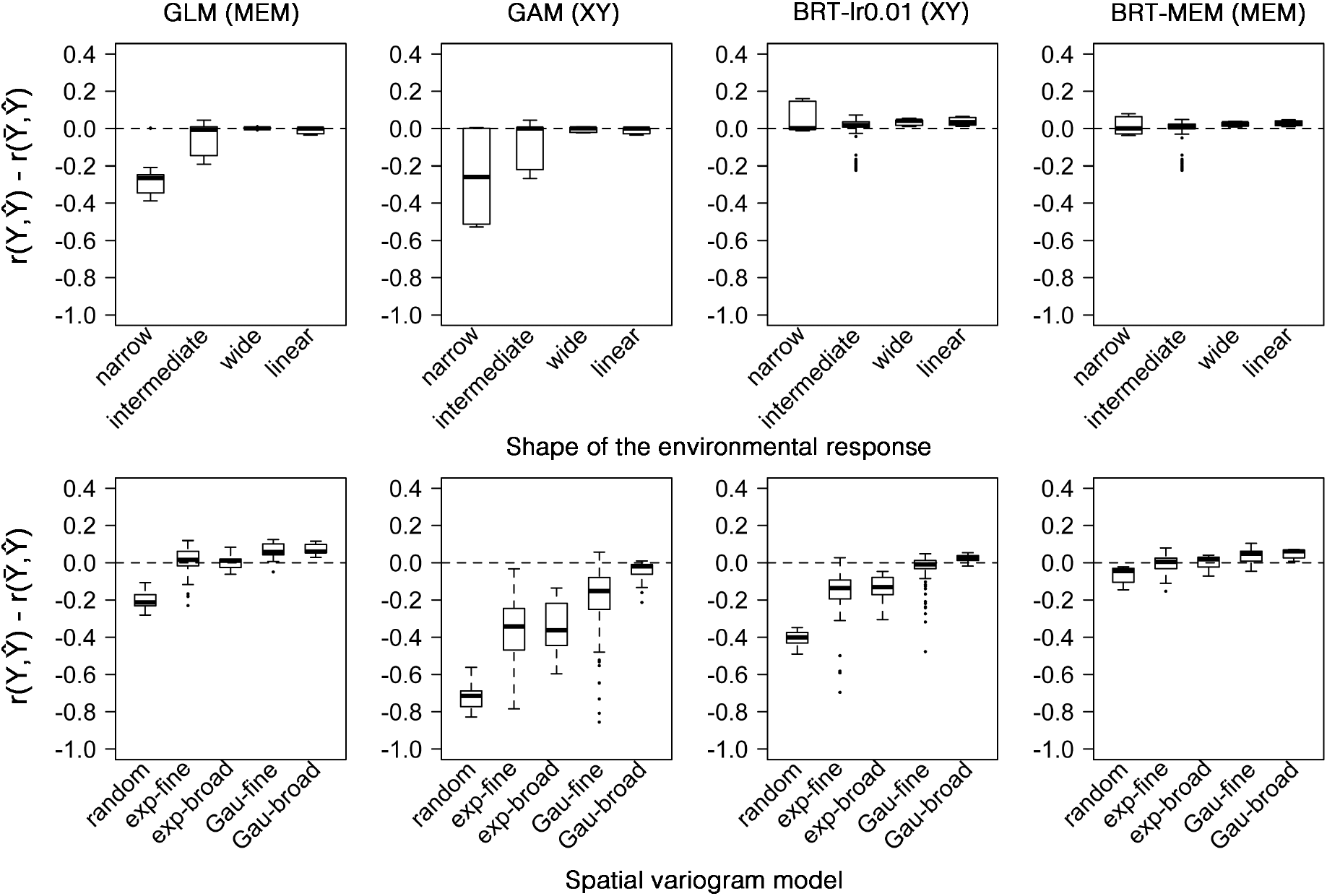
Fitting performance of different statistical models (GLM, GAM-k5, BRT-lr0.01) for the different environmental effects (a, b, c; unimodal with different niche breadths and linear; see also Appendix S1: Figure S1.1) and the different spatial effects (d, e, f; exponential and Gaussian variogram models with *A*=0.01, i.e., nearly random spatial structure, *A*=0.5*N* or *A*=*N*; the smoothness of the spatial structure increases to the right; see also Appendix S1: Figure S1.2). The results are pooled for all types of response data (i.e., normal, counts, and binary data). The performance is given by the correlation between the observed response values (*Y*) and the model predictions (*Ŷ*) minus the correlation between the simulated response values 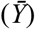 and the model predictions (*Ŷ*). The dashed line represents a perfect fit 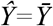, positive values indicate overfitting, and negative values indicate underfitting.

### Exercise 2. Variation partitioning

The variation partitioning performance, i.e. the difference between the estimated 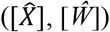and simulated ([X], [W]) fractions, varied considerably across methods, but was consistent across the different types of data (Fig. 3, 4). None of the methods perfectly recovered both the spatial and environmental fractions from the simulated reference variation partitioning. GLMs and BRT with MEMs (BRT-MEM) were the most consistent with the simulated variation partitioning (Fig. 3, 4). In contrast, the distance-based regression (MRM) severely underestimated both fractions of variation (Fig. 3). The results were qualitatively similar when using *R*^*2*^ or deviance-based pseudo-*R*^*2*^ metrics for partitioning the variation (Appendix S1; Fig. S1.7, S1.8).

**Figure 3.**
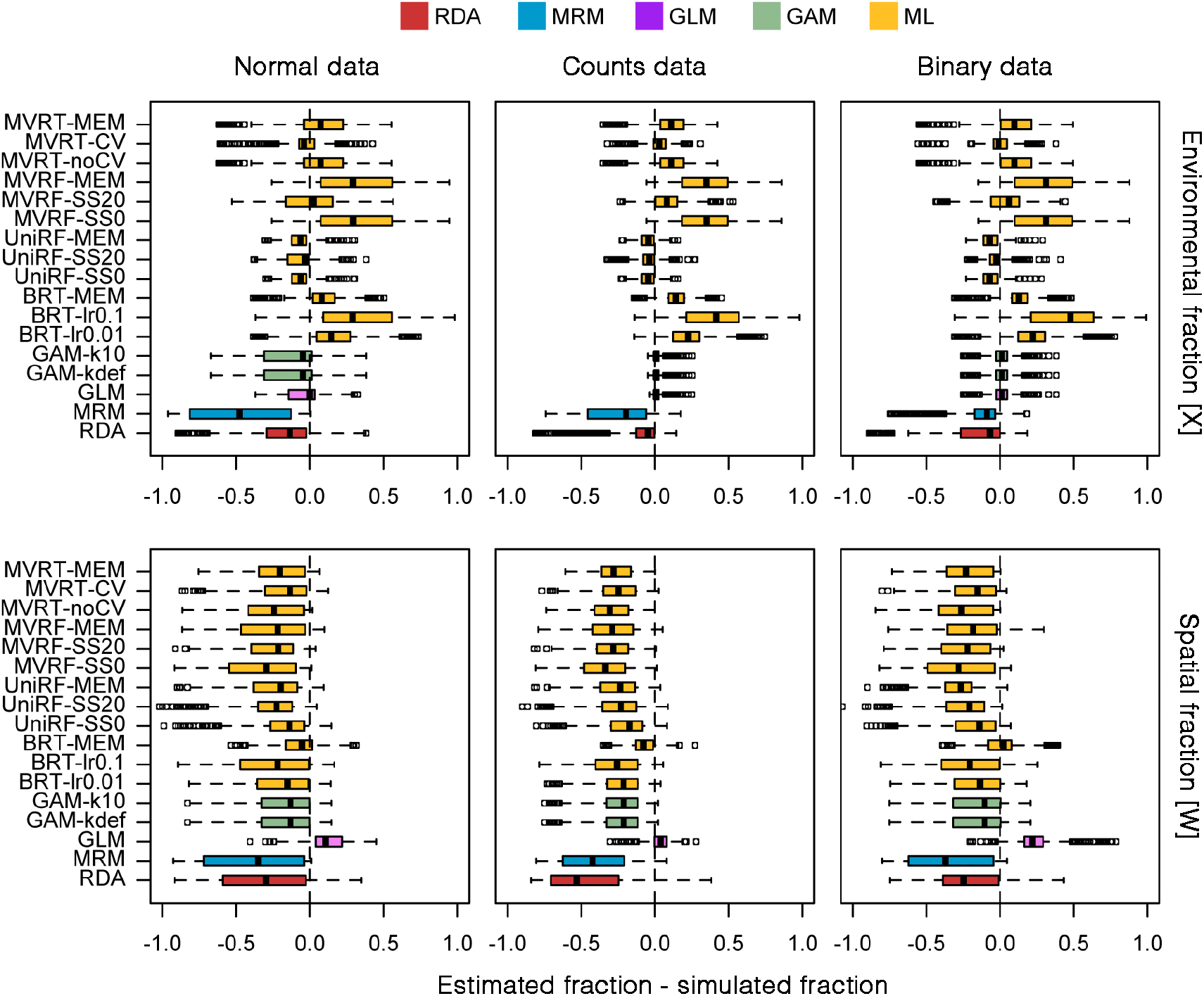
Variation partitioning performance of the different methods for normal, counts, and binary data across the different generated data. The performance is given by the difference between the estimated and simulated fractions of variation explained by the environment (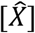 - [*X*]; upper panels) and space ([*Ŵ*] - [W]; lower panels). The dashed line represents equality (simulated = estimated), positive values indicate overestimation, and negative values indicate underestimation.

**Figure 4.**
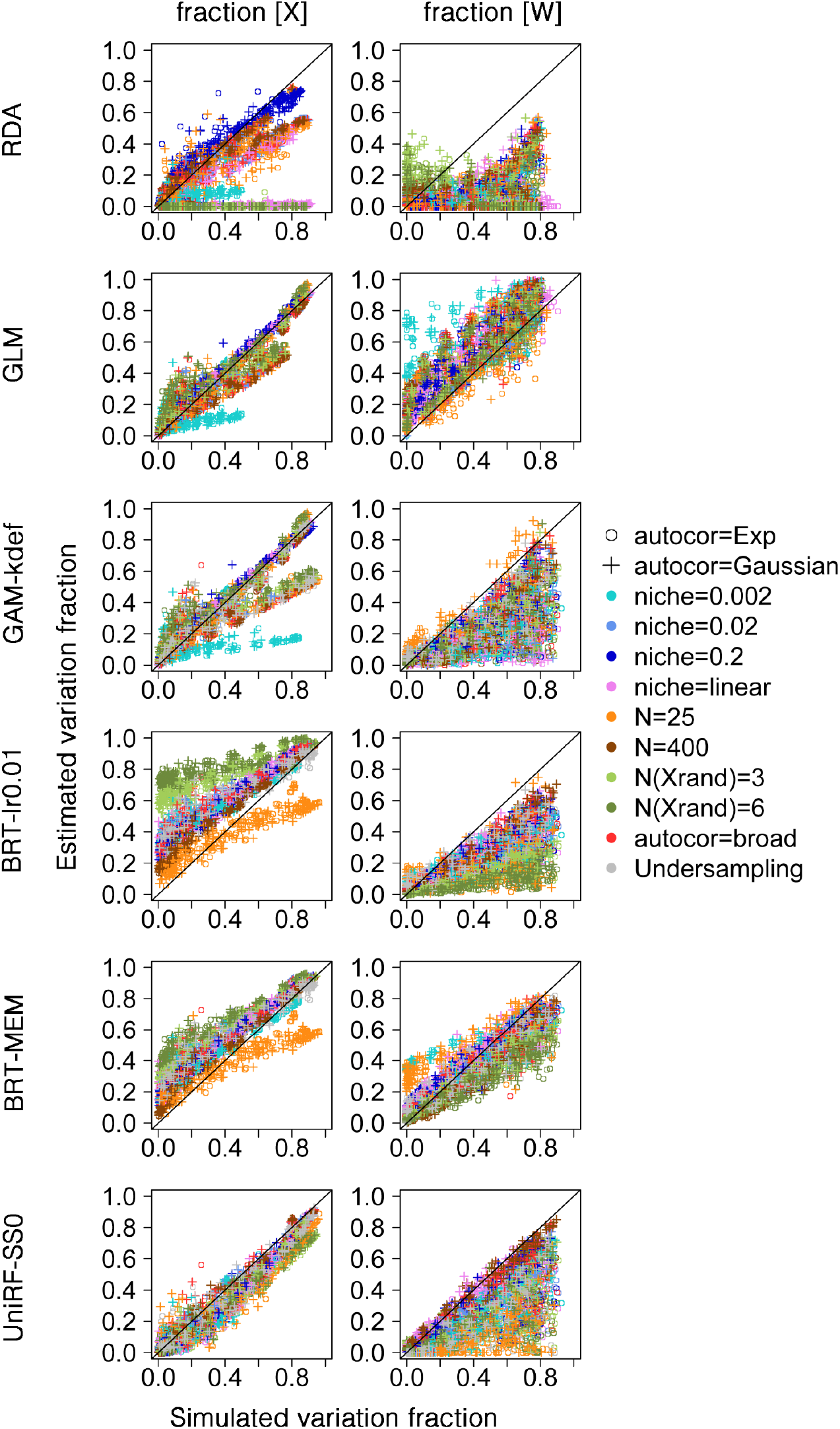
Plots of simulated vs. estimated fractions of variation explained by the environment ([*X*] vs.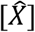; left panels) and space ([W] vs. ([*Ŵ*]; right panels) across different methods (different rows) and generated data (different colours). The black line represents the 1:1 line, where points should fall if the estimated fraction is equal to the simulated fraction.

For GLMs, the tendency to generally overfit the spatial component led to the overestimation of the spatial fraction of variation [*Ŵ*] (Fig. 3). The environmental fraction was generally correctly estimated, but was underestimated for the narrowest bell-shaped responses – variation that was instead fitted by the spatial component (Fig. 4). RDA generally largely underestimated the environmental and spatial fractions of variation – note, however, that the spatial fraction was systematically higher when using the classical *R*^*2*^ metric, although its trend did not generally follow that of the simulated data (Appendix S1; Fig. S1.7 and S1.8).

GAMs underestimated the spatial fraction of variation but generally correctly estimated the environmental fraction. Still, and even though we used splines to fit the environmental effect, the narrowest responses to the environment were also underestimated. Because we let the smoothing parameter to be optimized, we did not find differences between the number of dimensions of the basis functions (default or k=10).

The performance of machine learning differed to a large extent across its different methods, as well as between the variants of each method depending on the type of spatial variables used and the parameters controlling model smoothness. All ML methods tended to overestimate the environmental fraction of variation, except UniRF and MVRT with cross-validation (MVRT-CV) (Fig. 3). On the contrary, the spatial fraction was generally underestimated, though only slightly for BRT with MEMs (Fig. 3, 4). The less smoothened versions of BRT (BRT-lr0.1) and MVRF (MVRF-SS0) largely overestimated the environmental fraction, but had only minor effects on the estimation of the spatial fraction (Fig. 3). BRT with MEMs was the method that performed consistently better across all simulated scenarios (Fig. 4). However, BRT was sensitive to sample size and the number of environmental covariates, noticeably overestimating the environmental fraction when we added environmental covariates unrelated to the response.

### Empirical data

The choice of method had a clear influence on the results of the variation partitioning on empirical data (Fig. 5). Fitting GLM and MVRT with cross-validation (MVRT-CV) led to convergence problems for some data sets, likely because of the large number of spatial predictors relative to sample size in GLMs, and the too few observations for cross-validation in MVRT. Consistently with the simulation exercises, the environmental and spatial fractions in RDA were systematically underestimated relative to GAM and BRT. The results for GAM were more similar to BRT, in particular for the environmental fraction of variation. The results for BRT with spatial coordinates (BRT-lr0.01) and MEMs (BRT-MEM) as spatial predictors had also the same trend, but the spatial fraction was systematically higher in BRT-MEM.

**Figure 5.**
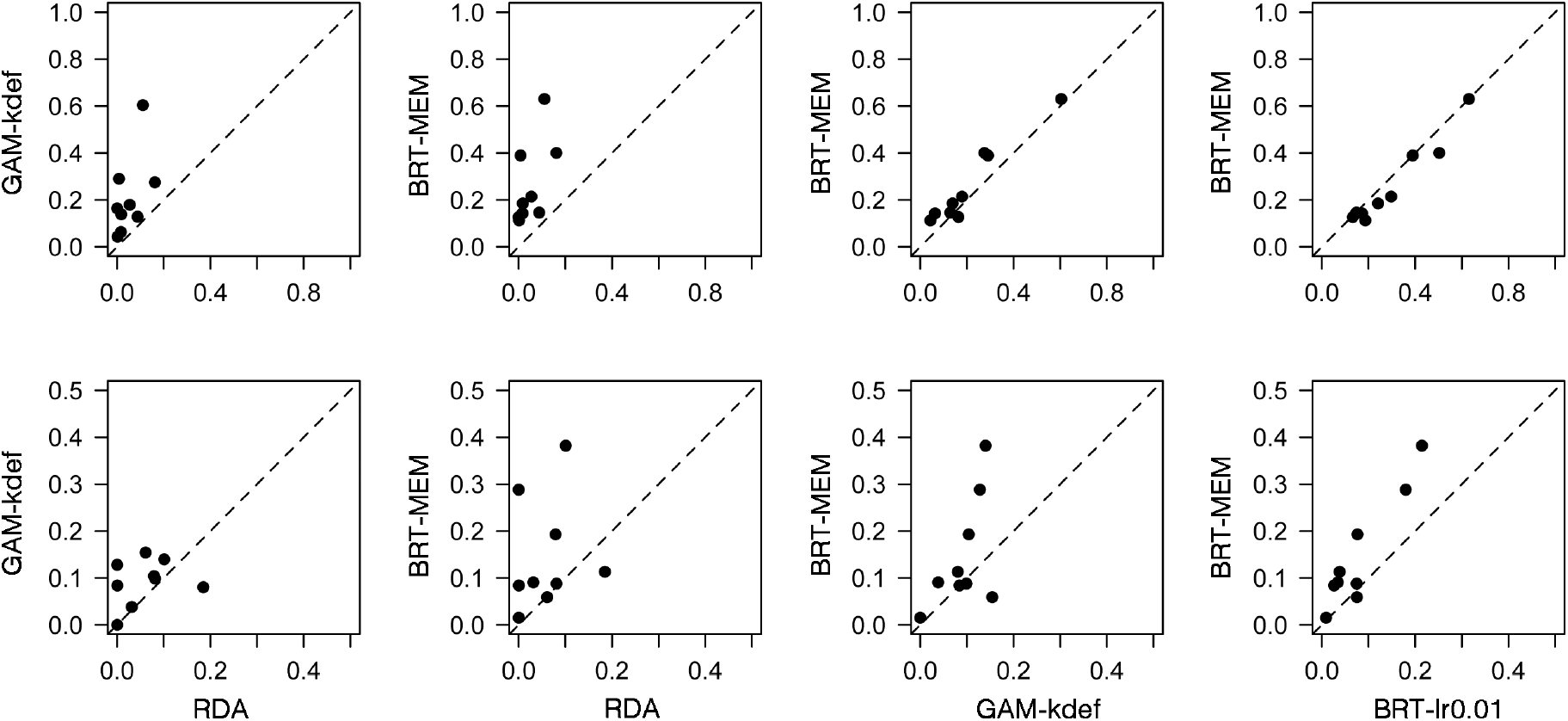
Cross-comparison of the estimated fractions of variation explained by the environment (upper panels) and space (lower panels) between RDA (as a widely used variation partitioning method for multivariate data), GAM, and a tree-based method (BRT; either with MEMs as spatial predictors or with spatial coordinates and a learning rate of 0.01) for the empirical datasets. The dashed line represents the 1:1 line. See Appendix S2 for references.

## Discussion

Our simulations show that it is challenging to recommend a single universal method for modelling spatial and environmental effects that can be used under any circumstances. All methods assessed here have advantages and drawbacks, and none of the methods provides a perfect model that always correctly fits both the environmental and spatial effects. GLMs and BRT with MEMs can provide more balanced fits that more consistently disentangle the spatial and environmental sources of variation in species abundance and occurrence. This highlights the potential of GLMs as an alternative to popular constrained ordination methods, and of BRT combined with powerful spatial predictors (MEMs) as a flexible, yet not widely deployed, method for ecologists aiming at disentangling environmental and spatial effects. GLMs are particularly useful when specific hypotheses about species-environment relationships are considered, as we can limit the scope of modelled responses to, for instance, linear and unimodal. In contrast, BRT allows us to fit any kind of response. The other methods can also be useful under certain situations.

GAMs are a good compromise, as modelled species-environment relationships can be limited to parametric response shapes (e.g. polynomials or splines with a limited number of basis functions, *k*) while spatial effects can be fitted using flexible splines on spatial coordinates. Because GAMs tend to fit smoother spatial surfaces in comparison to other methods, they might be more transferable, for example if the goal is to predict the response in unsampled sites. Note, however, that these models underestimate spatial effects when spatial autocorrelation acts over short distances. To allow GAMs to be flexible enough as to model wobblier effects, we can for example perform model selection according to *k* and smoothness of the splines (Wood 2007).

Tree-based ML methods have the advantage of being flexible and user-friendly, and are especially useful when we lack specific hypotheses about the functional forms of species-environment relationships, even though ML output can be interpreted and tested for effect significance (Lucas 2020). Furthermore, tree-based ML inherently models interactions between predictors, a desirable property when modelling complex ecological systems. However, it was challenging to parameterize ML algorithms that worked well for jointly fitting the environmental and spatial components. Simply using spatial coordinates to model space turned out to underfit most spatial structures, but forcing the model to be less smooth often led to overfitting of the environmental component. The use of MEMs in BRT models can offer a more balanced solution for jointly estimating environmental and spatial effects. However, this was not the case for other ML methods, for reasons that remain elusive. Thus, although ML showed potential to be used for variation partitioning purposes, we still need to learn more about how to fine-tune the ML parameters affecting spatial structure.

This brings us to the challenge of modelling spatial structures. Among the spatial structures that we considered, those generated with exponential variograms (Appendix S1: Figure S1.2) tended to be underfitted, whereas nearly random spatial structures tended to be overfitted by the methods that better fitted the former (Fig. 2). Therefore, it is critical to choose the type of spatial structure depending on the processes involved (e.g. dispersal limitation or connectivity) and goals of the study. MEMs are useful for interpretation purposes, for example to infer spatial scales of variation (Dray et al. 2012, Murakami and Griffith 2019). GAMs, on the other hand, can fit smooth spatial surfaces (e.g. Appendix S1: Figure S1.6), such as those simulated via a Gaussian variogram model (Appendix S1: Figure S1.2). And ML can offer flexible solutions, but *a priori* explorations, e.g., through simulations, might be needed in order to know how flexible the spatial structure needs to be. We also warn that increasing numbers of spatially autocorrelated environmental predictors generate spurious correlations (Chapman 2010), something that shall be accounted for when performing variation partitioning (Clappe et al. 2018 and the method therein). Cross-validation is a good solution, but it should be performed carefully in a spatial setting, because fitted spatial surfaces are hardly transferable in space, and training and validation data may not be spatially independent (Roberts et al. 2017). If the goal is to estimate the total fraction of variation explained by the environment independently of space (i.e., if spatial effects are not of interest), and if sample size is sufficient, then spatially blocked cross-validation is a good option (Roberts et al. 2017).

Our results show that there is margin for improvement. ML methods are diverse, and more methods can be tested and even developed specifically for the purposes outlined here (e.g. D’Amen et al. 2017, Nieto-Lugilde et al. 2018). Also, joint species distribution models (JSDM), which are based on generalized linear modelling, are becoming popular in community ecology (e.g. Ovaskainen et al. 2017), but their performance to model different spatial structures remains poorly explored. It is also possible to combine ML techniques with JSDMs, which offer a more flexible approach to model non-linear responses while using JSDM machinery (Harris 2015). Other possibilities include the use of autoregressive models (Dormann et al. 2007), though we are unware of their use for variation partitioning, the development of better cross-validation procedures to partition explained variation, and the development of *R*^*2*^ adjustment procedures, in particular for ML methods such as BRT, since the addition of irrelevant covariates in the models resulted in the overestimation of the environmental fraction of variation.

In conclusion, we provide a comprehensive assessment of different methods to use in ecology when we need to jointly model environmental and spatial effects. As the particular shapes of species responses to the environment are hard to hypothesize in the context of community or multi-species data, we introduce tree-based ML as a flexible method that can be widely used with both abundance and occurrence data. If *a priori* hypotheses about species-environment relationships are considered, GLM as a parametric method is a reasonable choice for variation partitioning. By choosing appropriate methods to model different responses to the environment, and different spatial structures at different spatial scales, with typical ecological data, such as species abundances and distributions, species diversity, and community composition, our recommendations apply to community ecology, biogeography, and macroecology studies.

## Acknowledgements

We thank Jonathan Chase and Pedro Peres-Neto for important discussions and comments. The work was supported by the German Centre for Integrative Biodiversity Research (iDiv) Halle-Jena-Leipzig funded by the German Research Foundation (FZT 118). DSV was supported by sDiv, the Synthesis Centre of iDiv. PK was supported by REES (Research Excellence in Environmental Sciences) grant from Faculty of Environmental Sciences, Czech University of Life Sciences in Prague.

