## Appendix S1 for "Disentangling spatial and environmental effects: flexible methods for community ecology and macroecology"

**Supporting Information**

Appendix S1: Additional results

Duarte S. Viana, Petr Keil, Alienor Jeliazkov

**Supplementary Tables and Figures**

**Table S1.** Summary of parameters of the data simulation.

| **Component** | **Parameter** | **Description** | **Value** |
| --- | --- | --- | --- |
| Initial conditions | *N* | Grid size (number of cells) and thus sample size of the simulated data (number of sites) | 25, 100, 400 |
|  | *J* | Number of species | 20 |
| Environment  (*X*) | *E* | Environmental variable (uniform distribution) | 0 - 1 |
|  | *autocor* | Variogram model | Exponential |
|  | *R* | Range of the variogram defining the "grain" of environmental autocorrelation | 0.5*N* |
|  | *β_i_* | Slope of linear response of species *i* | N(*μ=*10, *σ=*2) |
|  | *μ_i_* | Optimal environmental condition of species *i* | 0.05 - 0.95 |
|  | *2σ^2^* | Niche breadth (equal among species) | 0.002, 0.02, 0.2 |
|  | *N(X_rand_)* | Number of random environmental predictors (besides *E*) | 0, 3, 6 |
|  | *β_X_* | Weight of the environmental component | 0 - 1 |
| Space (*W*) | *autocor* | Variogram model | Exponential, Gaussian |
|  | *R* | Range of the variogram defining the "grain" of autocorrelation | 0.01*N,* 0.5*N, N* |
|  | *β_W_* | Weight of the spatial component | 0 - 1 |

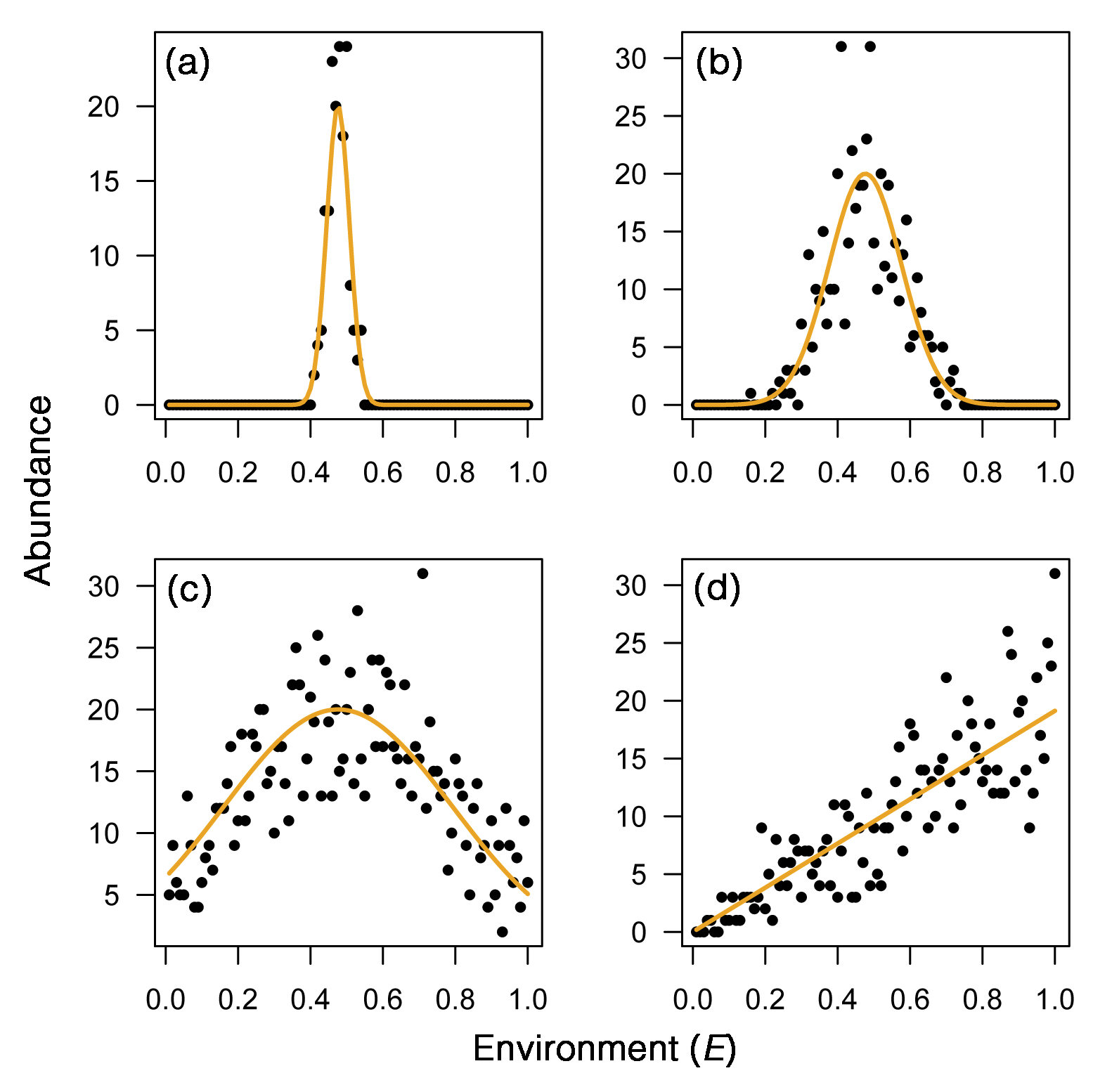

**Figure S.1.1.** Environmental effect (*X*): Gaussian-shaped responses (a-c) with increasing standard deviation (i.e., increasing "niche breadth") and linear response (d).

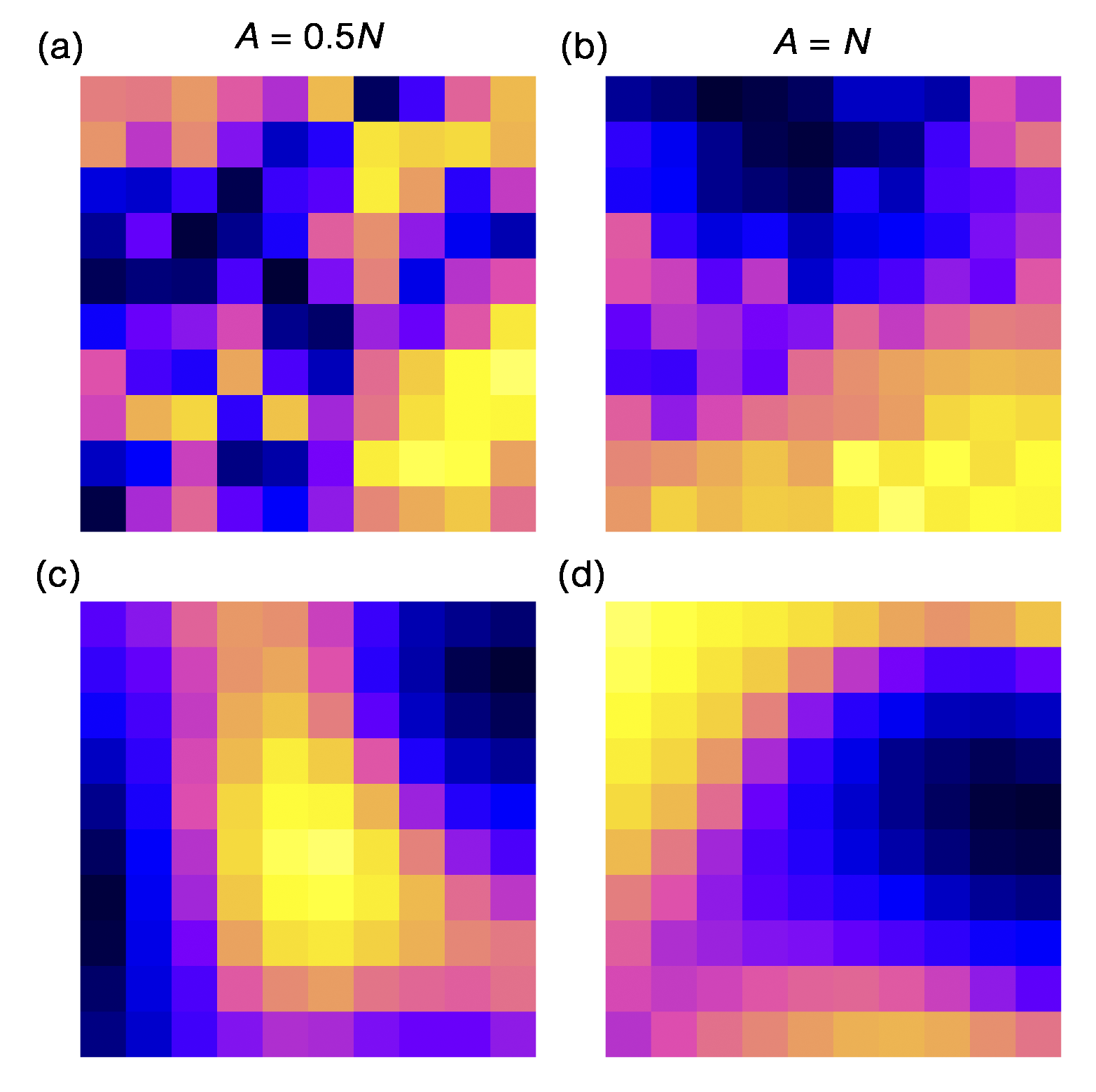

**Figure S.1.2.** Spatial effect (*W*): exponential variogram models with increasing range (*A*) parameter (a, b), and Gaussian variogram model with increasing range (*A*) parameter (c, d).

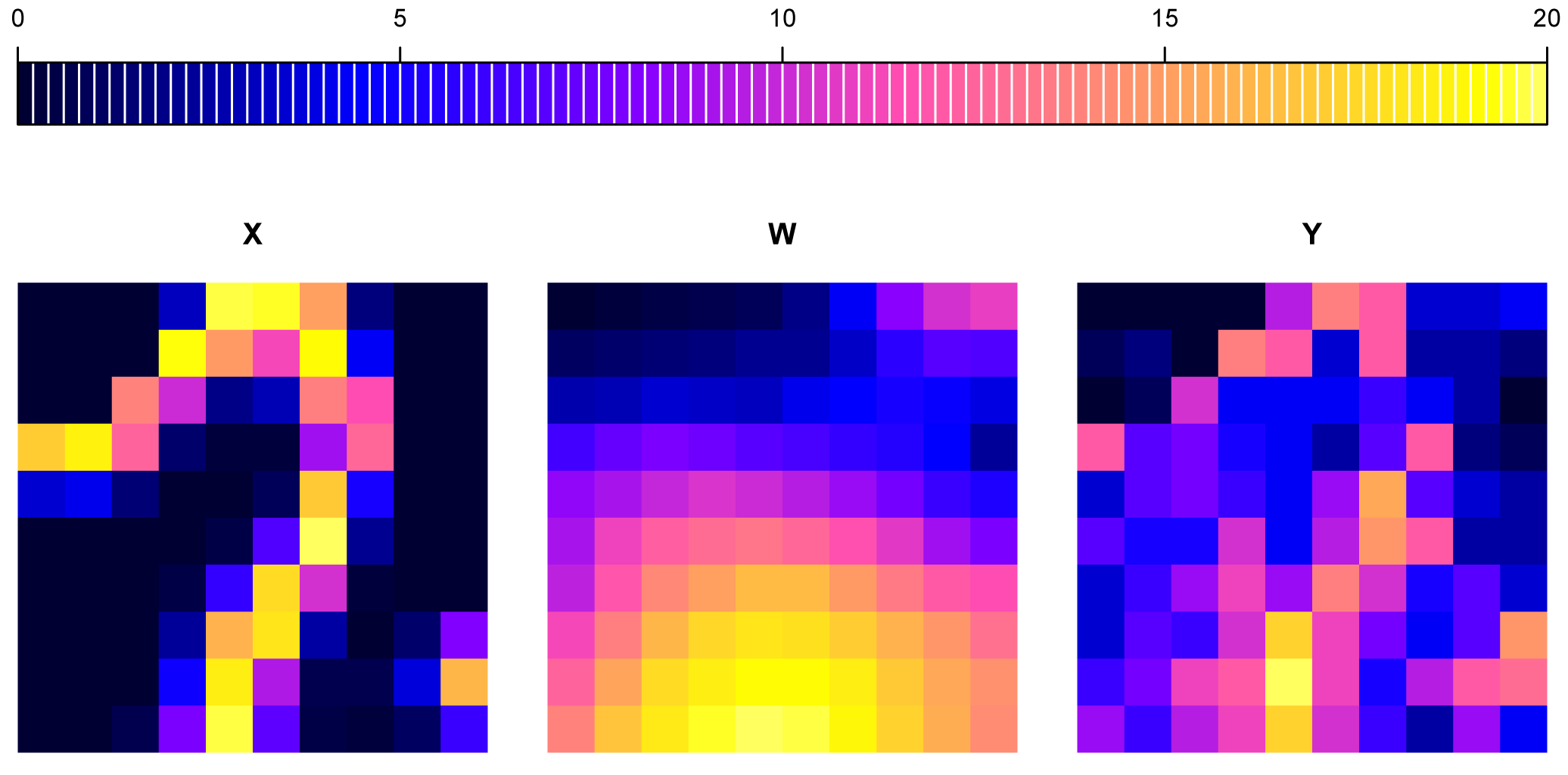

**Figure S1.3.** Spatial pattern of abundance of one species (from 0 to 20 individuals) of the environmental component (*X*), spatial component (*W*), and final additive abundance after a Poisson resampling (*Y*).

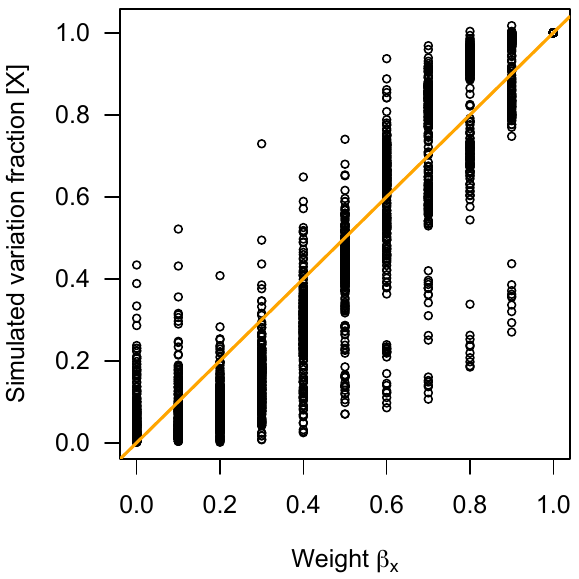

**Figure S1.4.** Relationship between the weights of the environmental component (*β_X_*) and the squared Pearson's correlation values between *X_i_* and *Y_i_* corresponding to the variation fraction explained by the simulated environmental effect. All the simulated data used in our study are represented here.

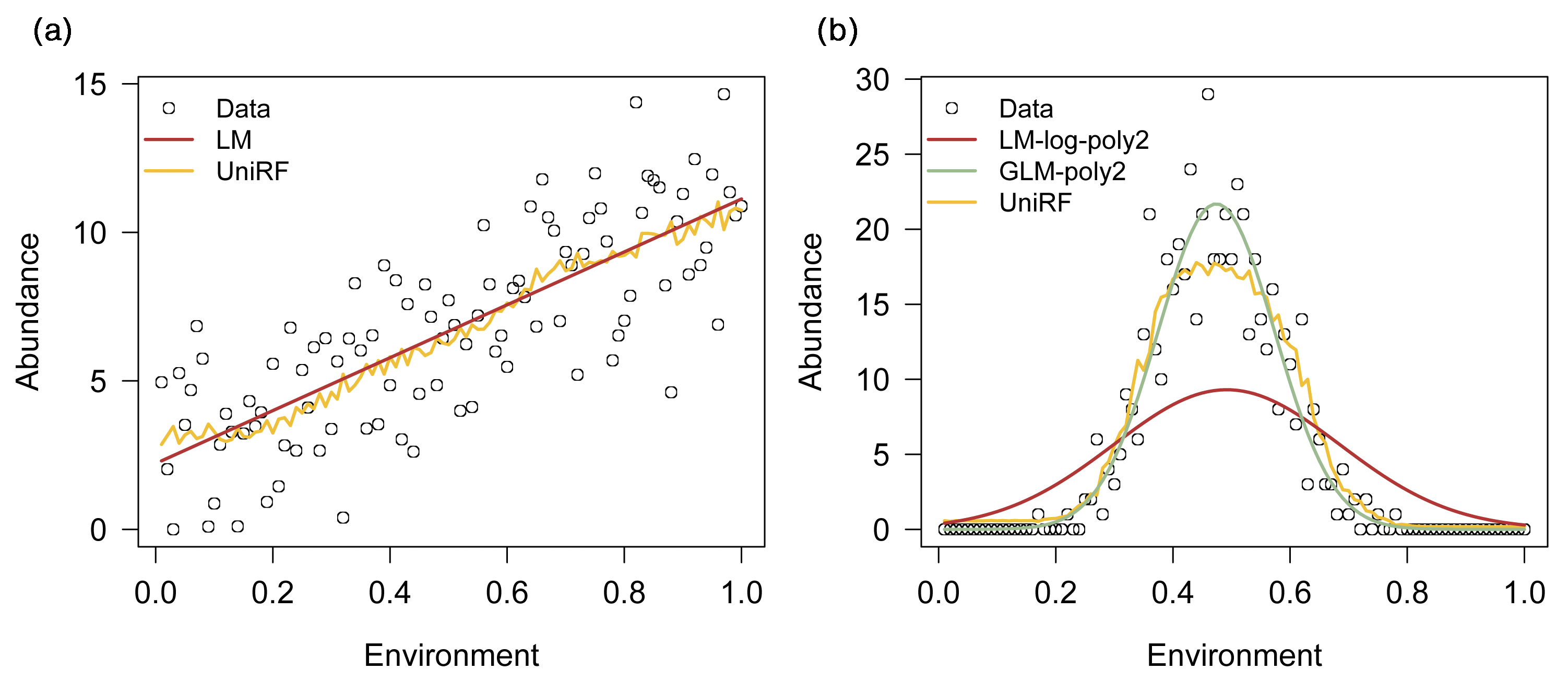

**Figure S1.5.** Example of models fitted to responses to the environment (*E*): a linear response (a) and a Gaussian-shaped response (b) of one species. For the linear response, a simple linear model (LM) and univariate random forest (UniRF) are fitted. For the Gaussian-shaped response (counts), a LM with log-transformed response data and a quadratic effect (LM-log-poly2), a Poisson generalised linear model (GLM) with a quadratic effect, and UniRF are fitted.

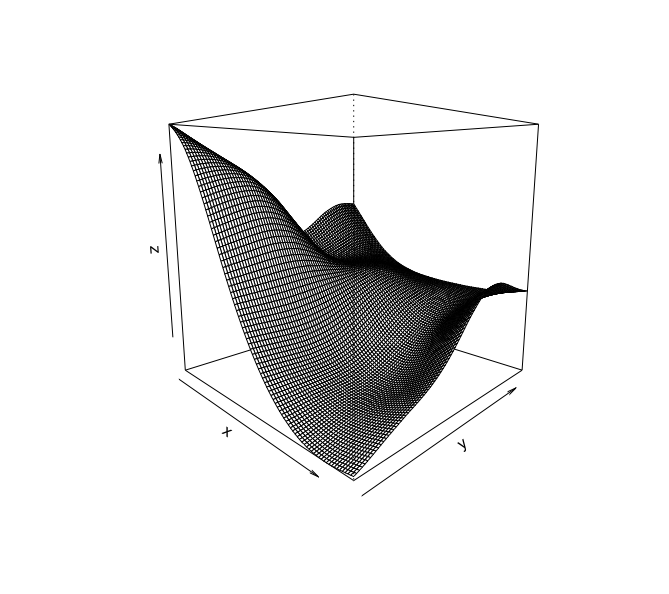

**Figure S1.6.** Example of non-linear spatial surface fitted to abundance data (z) by a spline through the geographic coordinates (x and y) in the GAM model.

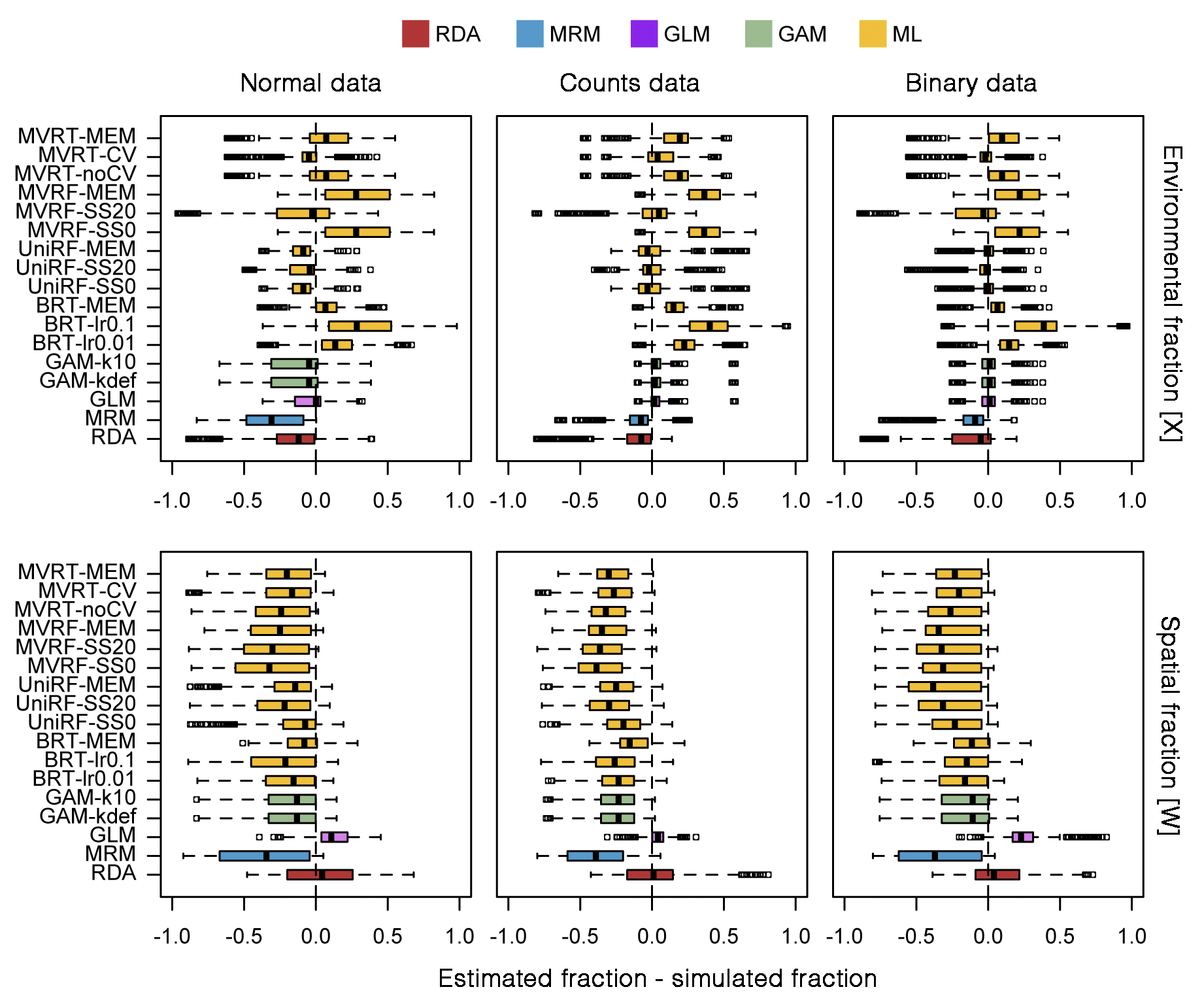

**Figure S1.7.** Variation partitioning performance of the different methods for normal, counts, and binary data across the different generated data. The performance is given by the difference between the estimated and simulated fractions of variation explained by the environment ($[\hat{X}]$ - [*X*]; upper panels) and space ([*Ŵ*] - [W]; lower panels). Here, the variation fractions are estimated using either the classic R^2^ or a deviance-based pseudo-R^2^ instead of r^2^ as in the main text. The dashed line represents equality (simulated = estimated), positive values indicate overestimation, and negative values indicate underestimation.

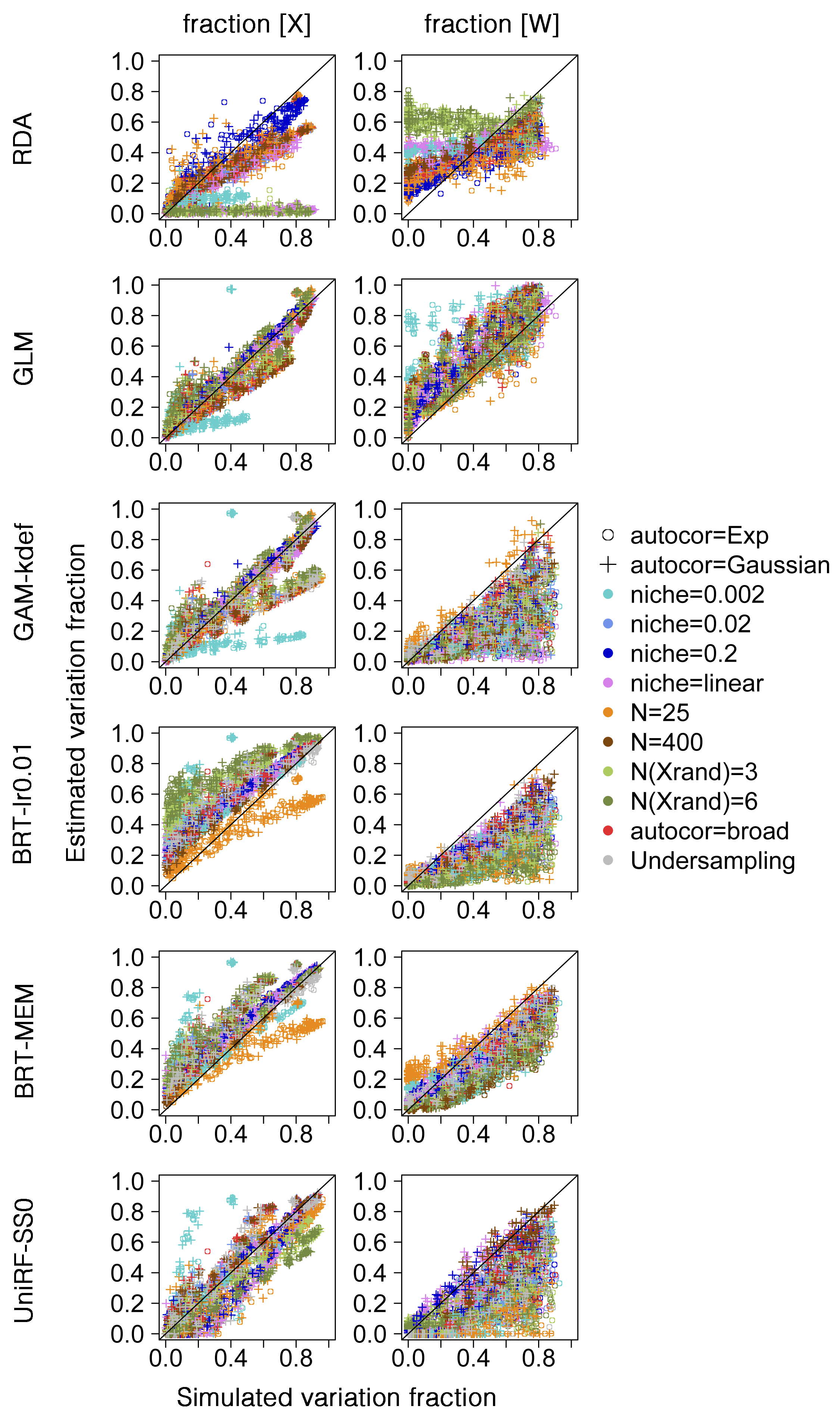

**Figure S1.8.** Plots of simulated vs. estimated fractions of variation explained by the environment ([*X*] vs. $[\hat{X}]$; left panels) and space ([W] vs. [*Ŵ*]; right panels) across different methods (different rows) and generated data (different colours). Here, the variation fractions are estimated using either the classic R^2^ or a deviance-based pseudo-R^2^ instead of r^2^ as in the main text. The black line represents the 1:1 line, where points should fall if the estimated fraction is equal to the simulated fraction.
