## Appendix S2 for "Disentangling spatial and environmental effects: flexible methods for community ecology and macroecology"

**Supporting Information**

Appendix S2

Duarte S. Viana, Petr Keil, Alienor Jeliazkov

### Table S2. Dataset sources and citation references

| **Dataset name** | **Taxonomic group** | **Source** | **References** |
| --- | --- | --- | --- |
| Bagaria2012 | plants | Publication Supp. Mat. | Bagaria, G., J. Pino, F. Rodà, & M. Guardiola, 2012. Species traits weakly involved in plant responses to landscape properties in Mediterranean grasslands. Journal of Vegetation Science 23: 432–442. |
| Borcard1992 | mites | R {ade4} | Borcard, D., Legendre, P., and Drapeau, P. (1992) Partialling out the spatial component of ecological variation. Ecology,73, 1045–1055. |
| Condit2002 | trees | R {vegan} | Condit, R, Pitman, N, Leigh, E.G., Chave, J., Terborgh, J., Foster, R.B., Nuñez, P., Aguilar, S., Valencia, R., Villa, G., Muller-Landau, H.C., Losos, E. & Hubbell, S.P. (2002). Beta-diversity in tropical forest trees. Science 295, 666–669.  Svenning J.-C., Kinner D.A., Stallard R.F., Engelbrecht B.M.J. & Wright S.J. (2004) Ecological determinism in plant community structure across a tropical forets landscape. Ecology 85, 2526–2538. |
| Doledec1996 | birds | R {ade4} + https://pbil.univ-lyon1.fr/R/pdf/ter7.pdf | Dolédec, S., D. Chessel, C. J. F. ter Braak, & S. Champely, 1996. Matching species traits to environmental variables: a new three-table ordination method. Environmental and Ecological Statistics 3: 143–166. |
| Gibb2015 | spiders | Publication | Gibb, H., D. Muscat, M. R. Binns, C. J. Silvey, R. A. Peters, D. I. Warton, & N. R. Andrew, 2015. Responses of foliage-living spider assemblage composition and traits to a climatic gradient in Themeda grasslands: Spider Traits and Climatic Gradients. Austral Ecology 40: 225–237. |
| Jeliazkov2014 | amphibians | https://idata.idiv.de/ddm/data/showdata/286 | Jeliazkov A., Chiron F., Garnier J., Besnard A., Silvestre M. & Jiguet F. (2014) Level-dependence of the relationships between amphibian biodiversity and environment in pond systems within an intensive agricultural landscape. Hydrobiologia 723, 7–23.  Jeliazkov & the CESTES consortium (2019) A global database for metaCommunity Ecology: Species, Traits, Environment and Space - version 1.0 (CESTES v1.0). |
| Pavoine2011 | plants | R {ade4} | de Bélair, Gérard and Bencheikh-Lehocine, Mahmoud (1987) Composition et déterminisme de la végétation d'une plaine côtière marécageuse : La Mafragh (Annaba, Algérie). Bulletin d'Ecologie, 18(4), 393–407. Pavoine, S., Vela, E., Gachet, S., de Bélair, G., & Bonsall, M. B. (2011). Linking patterns in phylogeny, traits, abiotic variables and space: a novel approach to linking environmental filtering and plant community assembly: Multiple data in community organization. Journal of Ecology, 99(1), 165–175. doi:10.1111/j.1365-2745.2010.01743.x |
| Ribera2001 | beetles | Publication Supp. Mat. | Ribera, I., S. Dolédec, I. S. Downie, & G. N. Foster, 2001. Effect of Land Disturbance and Stress on Species Traits of Ground Beetle Assemblages. Ecology 82: 1112–1129. |
| Verneaux1973 | fishes | R {ade4} | Verneaux, J. (1973) Cours d'eau de Franche-Comté (Massif du Jura). Recherches écologiques sur le réseau hydrographique du Doubs. Essai de biotypologie. Thèse d'état, Besançon. 1–257. |
